# A year in the life of the Eastern Mediterranean: Monthly dynamics of phytoplankton and bacterioplankton in an ultra-oligotrophic sea

**DOI:** 10.1101/2021.03.24.436734

**Authors:** Tom Reich, Tal Ben-Ezra, Natalya Belkin, Anat Tsemel, Dikla Aharonovich, Dalit Roth-Rosenberg, Shira Givati, Or Bialik, Barak Herut, Ilana Berman-Frank, Miguel Frada, Michael D. Krom, Yoav Lehahn, Eyal Rahav, Daniel Sher

## Abstract

The Eastern Mediterranean Sea (EMS) is a poorly studied ultra-oligotrophic marine environment, dominated by small-size phyto- and bacterioplankton. Here, we describe the dynamics of a single annual cycle (2018-19) of phyto- and bacterioplankton (abundances, pigments and productivity) in relation to the physical and chemical conditions in the photic water column at an offshore EMS site (Station THEMO-2, ∼1,500m depth, 50km offshore). We show that phytoplankton biomass (as chlorophyll a), primary and bacterial productivity differed between the mixed winter (January-April) and the thermally stratified (May-December) periods. *Prochlorococcus* and *Synechococcus* numerically dominated the picophytoplankton populations, with each clade revealing different temporal and depth changes indicative to them, while pico-eukaryotes (primarily haptophytes) were less abundant, yet likely contributed significant biomass. Estimated primary productivity (∼32 gC m^-2^ y^-1^) was lower compared with other well-studied oligotrophic locations, including the north Atlantic and Pacific (BATS and HOT observatories), the western Mediterranean (DYFAMED observatory) and the Red Sea, and was on-par with the ultra-oligotrophic South Pacific Gyre. In contrast, integrated bacterial production (∼11 gC m^-2^ y^-1^) was similar to other oligotrophic locations. Phytoplankton seasonal dynamics were similar to those at BATS and the Red Sea, suggesting an observable effect of winter mixing in this ultra-oligotrophic location. These results highlight the ultra-oligotrophic conditions in the EMS and provide, for the first time in this region, a full-year baseline and context to ocean observatories in the region.

**Highlights:**

- Bacterioplankton dynamics were assessed monthly in the Eastern Mediterranean Sea
- Small-sized picophytoplankton numerically dominated the phytoplankton community
- Seasonal phytoplankton dynamics are similar to BATS and Red Sea, but not to HOT
- Annual primary productivity is among the lowest in the world’s oceans
- Bacterial to primary production ratio is higher than most oligotrophic seas

## 1 Introduction

Convective mixing of the water column is one of the main mechanisms responsible for transport of nutrients to the photic zone of oligotrophic oceans and seas, often resulting in increased phytoplankton biomass and activity (Behrenfeld, 2010). This process usually occurs during wintertime upon the progressive cooling of the sea surface, although other mechanisms may also deliver nutrients to the mixed layer such as physical upwelling along shorelines, frontal systems and gyres (Anabalón et al., 2016), nutrient runoff from rivers (Jickells, 1998), and atmospheric deposition (Guieu et al., 2014). Stratification is established during springtime, nutrients gradually become depleted, which, together with other processes such as increased predation, lead to a decline in algal biomass and productivity (Behrenfeld and Boss, 2014). This cycle has been extensively studied over decades in the major oligotrophic gyres (e.g. HOT and BATS) (Steinberg et al., 2001), as well in other locations including the Western Mediterranean and Red Seas (Genin et al., 2018; Marty et al., 2002; Marty and Chiavérini, 2010), and is considered a fundamental driving force of marine ecosystem structure and function.

The offshore water of the Eastern Mediterranean Sea are considered ultra-oligotrophic (Berman-Frank and Rahav, 2012; Siokou-Frangou et al., 2010). The oligotrophic nature of this system, and especially of the Eastern Mediterranean Sea (EMS), is mainly driven by its general anti-estuarine circulation (Pinardi and Masetti, 2000). Additionally, the relatively stable density stratification throughout most of the year (winter mixing rarely exceeds the 200 m depth, (D’Ortenzio et al., 2005) has been suggested to result in a very low supply of deep water nutrients to the euphotic zone (Hazan et al., 2018). Finally, modern riverine inputs into the EMS are extremely low, especially since the Nile was dammed in the 1960s (Krom et al., 2014). The abovementioned conditions makes the EMS, and specifically the easternmost Levantine basin, among the warmest, saltiest and least productive waters in the world (Ozer et al., 2017).

Despite the importance of the EMS coastline in providing ecosystem services to over 70 million people e.g. (Peled et al., 2018), it lacks the continued high resolution records available from other oligotrophic regions such as the North Atlantic (the Bermuda Atlantic Time Series, BATS), North Pacific (The Hawaii Ocean Time-Series station ALOHA, HOT), the Western Mediterranean e.g. station DYFAMED (Marty et al., 2002; Marty and Chiavérini, 2010) and, to a lesser extent, the Red Sea (Shaked and Genin, 2017). Specifically, detailed phytoplankton and bacterioplankton time-series are missing, and are important in order to both understand the current system and the ways it may be impacted by local, regional and global change (e.g. (Marty and Chiavérini, 2010). Thus, studying the dynamics of the microorganisms at the base of the marine food-web over time are of great ecological importance.

Studies from the early 1980’s showed a clear seasonal cycle of chlorophyll a (a proxy of algal biomass) and primary production in surface waters of both neritic and pelagic locations, with the changes attributed primarily to picophytoplankton (Azov, 1986; Berman et al., 1984; Kimor and Wood, 1975; Yacobi et al., 1995). More recent studies of the EMS show similar seasonal trends using remote-sensing of surface chlorophyll a (Rosenberg et al., 2020) and in measurements of cell numbers, production and dinitrogen fixation in coastal waters (Rahav et al., 2018; Raveh et al., 2015). However, to the best of our knowledge, no detailed temporal (monthly) measurements have been presented of microbial processes and phytoplankton community structure at the offshore EMS waters. In this study, we followed the composition and activity of phytoplankton and bacterioplankton from a pelagic location (station THEMO-2) in the offshore EMS over a full year at monthly temporal resolution, and at high depth resolution (∼20 samples, ∼12 of them across the photic zone). These measurements were performed as part of the SoMMoS (<u>So</u>utheastern <u>M</u>editerranean <u>Mo</u>nthly cruise <u>S</u>eries) campaign, which compared an open-ocean station with one at the edge of the continental shelf (Figure 1A). Additional studies from this cruise series will focus on the carbonate system (Juntao et al. *in prep*), nutrient dynamics (Ben Ezra et al., 2021), coccolithophore dynamics (Keuter et al. *in prep*) and a detailed comparison of the offshore and coastal stations (Krom et al. *in prep*). Together, these studies provide a baseline allowing improved interpretation for future research from the two long-term ocean observatories recently established in the EMS: the DeepLev (Katz et al., 2020) and THEMO (Diamant et al., 2020), as well as comparison to long-term monitoring activities (Ozer et al., 2017; Rahav et al., 2019; Sisma-Ventura et al., 2021).

**Figure 1:**
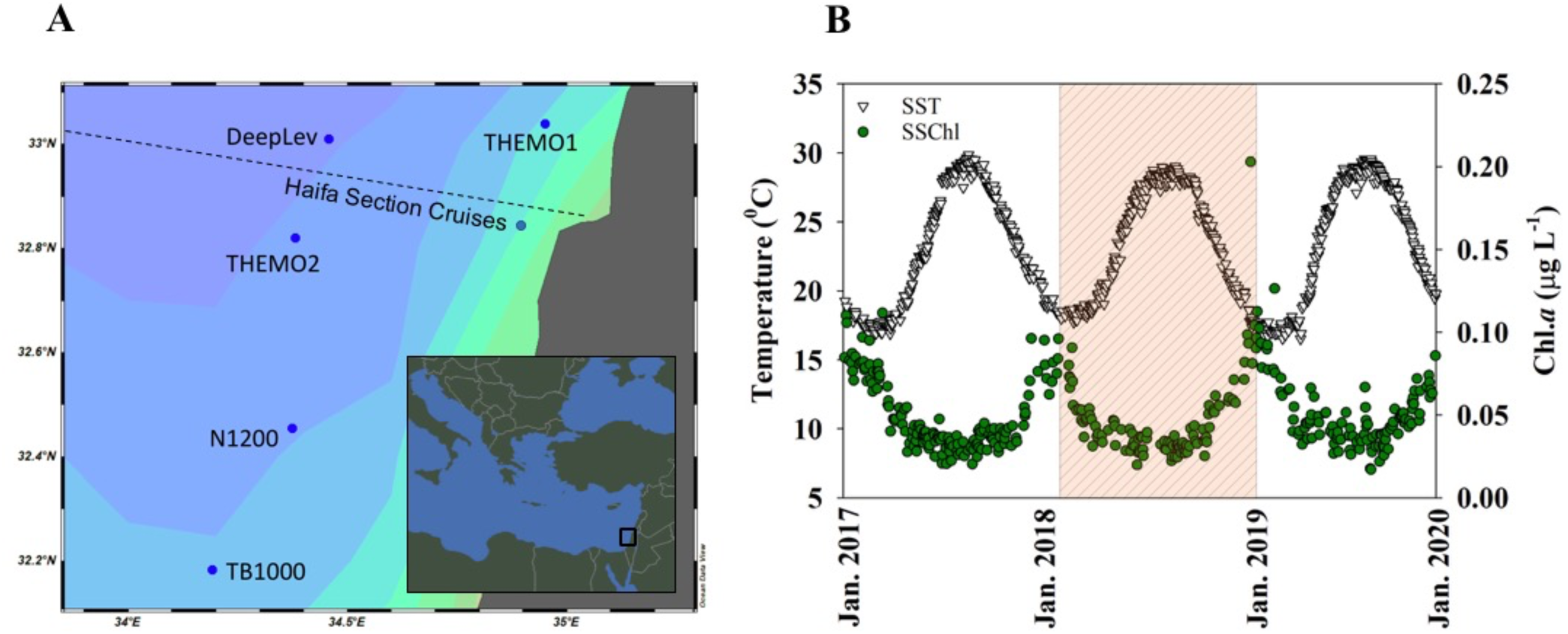
**Overview of the SoMMoS sampling locations context.** A) A map of the Levantine Basin (EMS) showing the location of the THEMO-2 sampling site (the focus of this study), as well as of additional observatories and locations of long-term studies discussed here: THEMO-1, DeepLev (Katz et al., 2020), N-1200 (Rosenberg et al., 2020), TB1000 (Yogev et al., 2011) and the Haifa section transect cruises (Ozer et al., 2017). At the bottom right corner an inset of the EMS with a black frame indicating the study site. B) Changes in satellite-derived sea surface temperature (orange) and sea surface chlorophyll a (green) from the 1 km^2^ region surrounding THEMO-2 between 2017-2020. Data were extracted from the Copernicus Marine Environment Monitoring Service (CMEMS, https://marine.copernicus.eu/). The period of this study is shaded.

## 2 Materials and Methods

### 2.1 Cruises and sample collection

Water samples were collected as part of the SoMMoS project during twelve cruises, from February 2018 until January 2019. Each cruise took samples at two stations: THEMO2, an open ocean station at a depth of ∼1500 m (∼50 Km from the coast, 32.820 N, 34.380 E), and THEMO1 which is positioned at the edge of the continental shelf at ∼125m depth (∼10 Km from the coast, 33.040 N, 34.950 E). THEMO2 was sampled from 12:00-22:00 (local time), while THEMO1 was sampled during the night, typically 00:00-02:00 (local time). Here, we will present information from data collected at the offshore THEMO2 station. Samples were collected using a 12-bottle rosette with 8 L Niskin bottles. Samples were collected at 20-24 depths across the entire water column (11-12 bottles between the surface and 200 m, which we define here as the photic zone). Sampling depths were selected based on real-time data of Conductivity, Temperature, Depth (CTD) profiler (Seabird 19 Plus) from the down-cast before each sample collection in the up-cast, an oxygen optode (on some cruises) and a fluorescence meter (Turner designs, Cyclops-7). The continuous data was processed using the Sea-Bird data conversion software, and minimized using bin averaging. One bin of data lines was defined as a change of 1 decibar (db) between each bin. The first two meters of measurements were compiled together to account for sensors and CTD pump adjustment time and rosette depth while at sea surface. Each cast allocated water for analysis in the following order to give priority to the more ‘sensitive’ parameters: dissolved methane, pH, alkalinity, dissolved inorganic carbon, inorganic nutrients, total and dissolved organic carbon, cell count, primary and bacterial production, DNA and algal pigment markers. Pre-filtered inorganic nutrient samples were analyzed fresh (unfrozen) the day after the cruise, using a SEAL AA-3 autoanalyzer system, and are described in detail elsewhere (Ben-Ezra et al., 2021). A summary of all of the currently-available measurements can be found in Supplementary Table S1, and the BCO-DMO (acronym: SoMMoS) and ISRAMAR (https://isramar.ocean.org.il/isramar2009/) databases. Mixed layer depth (MLD) was calculated using a temperature difference of Δ0.3 °C (Mena et al., 2019). Calculations based on a density difference of Δ0.15 kg/m^3^ yielded similar results. During several months (February-April 2018 and January 2019), the density plots revealed a progressive increase in density without a clear pycnocline but with multiple “bumps”, indicative of water column instability down to below 200 m (Supplementary Figure S1). At these times, the MLD calculations based on a defined difference in temperature or salinity from the surface preclude a robust estimate of the mixed layer depth, as they may underestimate the actual values. Based on these calculations, we divide the study period into ‘a generally mixed period’ during January-April (winter/spring), and a ‘stratified period’ during May-December (summer/autumn, Table 1). Estimates based on the vertical distribution of inorganic nutrient concentrations suggest the mixing period may have begun as early as November 2018 (Ben-Ezra et al., 2021). We note that the monthly sampling resolution likely to ‘miss’ short-lived deep mixing events, as can be observed from mooring operations (e.g. (Gunn et al., 2020).

**Table 1:**
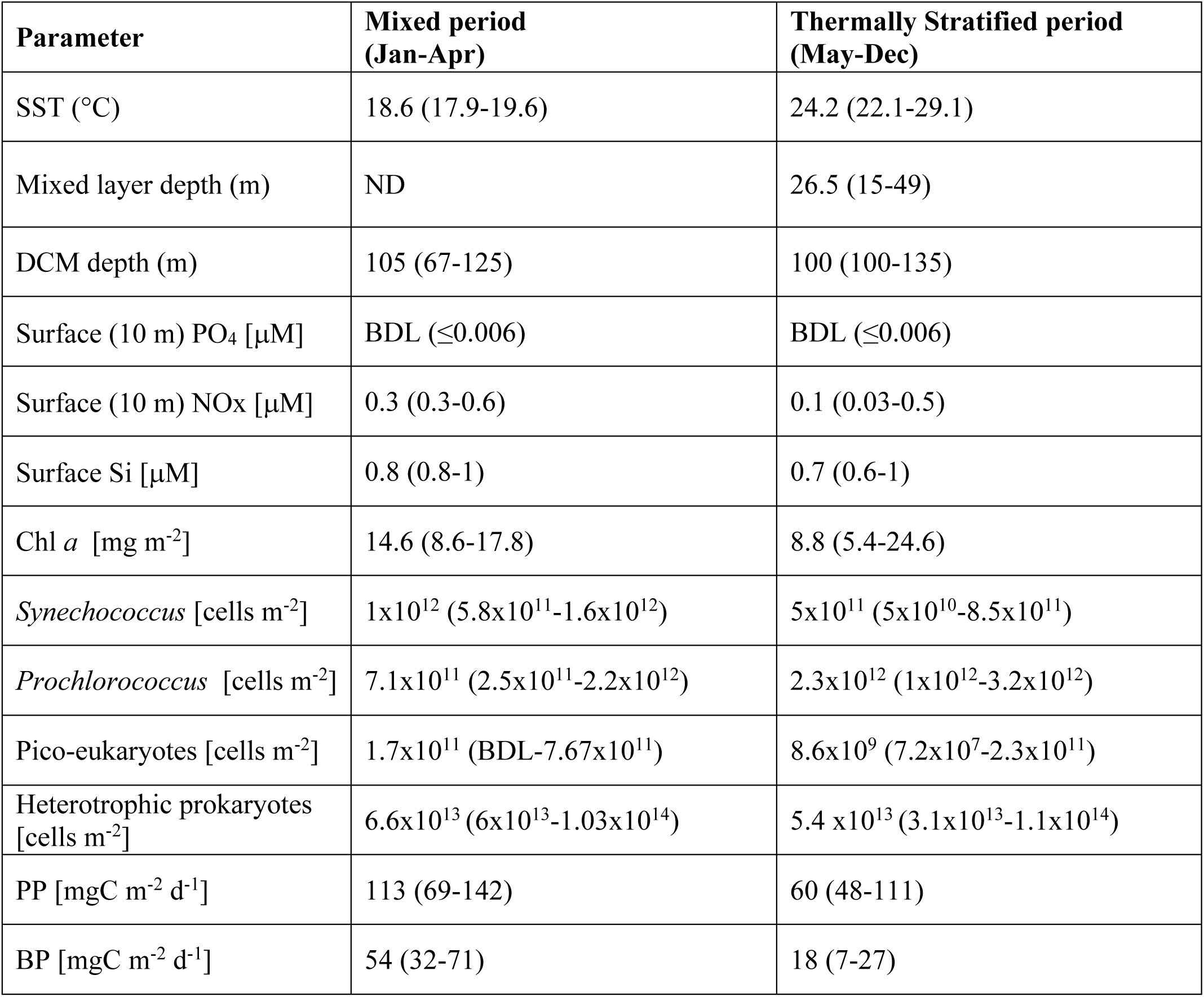
Median and range of major measured oceanographic parameters during the mixed and stratified periods. BDL – Below detection limit. ND – not determined (see materials and methods and Supplementary Figure S1 for details). Depth-integrated values are from 0-200m.

| Parameter | Mixed period<br>(Jan-Apr) | Thermally Stratified period<br>(May-Dec) |
| --- | --- | --- |
| SST (°C) | 18.6 (17.9-19.6) | 24.2 (22.1-29.1) |
| Mixed layer depth (m) | ND | 26.5 (15-49) |
| DCM depth (m) | 105 (67-125) | 100 (100-135) |
| Surface (10 m) PO<sub>4</sub> [μM] | BDL (≤0.006) | BDL (≤0.006) |
| Surface (10 m) NO<sub>x</sub> [μM] | 0.3 (0.3-0.6) | 0.1 (0.03-0.5) |
| Surface Si [μM] | 0.8 (0.8-1) | 0.7 (0.6-1) |
| Chl *a* [mg m<sup>-2</sup>] | 14.6 (8.6-17.8) | 8.8 (5.4-24.6) |
| *Synechococcus* [cells m<sup>-2</sup>] | 1x10<sup>12</sup> (5.8x10<sup>11</sup>-1.6x10<sup>12</sup>) | 5x10<sup>11</sup> (5x10<sup>10</sup>-8.5x10<sup>11</sup>) |
| *Prochlorococcus* [cells m<sup>-2</sup>] | 7.1x10<sup>11</sup> (2.5x10<sup>11</sup>-2.2x10<sup>12</sup>) | 2.3x10<sup>12</sup> (1x10<sup>12</sup>-3.2x10<sup>12</sup>) |
| Pico-eukaryotes [cells m<sup>-2</sup>] | 1.7x10<sup>11</sup> (BDL-7.67x10<sup>11</sup>) | 8.6x10<sup>9</sup> (7.2x10<sup>7</sup>-2.3x10<sup>11</sup>) |
| Heterotrophic prokaryotes<br>[cells m<sup>-2</sup>] | 6.6x10<sup>13</sup> (6x10<sup>13</sup>-1.03x10<sup>14</sup>) | 5.4 x10<sup>13</sup> (3.1x10<sup>13</sup>-1.1x10<sup>14</sup>) |
| PP [mgC m<sup>-2</sup> d<sup>-1</sup>] | 113 (69-142) | 60 (48-111) |
| BP [mgC m<sup>-2</sup> d<sup>-1</sup>] | 54 (32-71) | 18 (7-27) |

### 2.2 Bacterial and primary productivity

Heterotrophic prokaryotic productivity (hereafter referred to as bacterial productivity, BP) was estimated using the ^3^H-leucine incorporation method (Simon and Azam, 1989). Triplicate 1.7 ml of ocean water were taken from each sampled depth and incubated with a 7:1 mixture of ‘cold’ leucine and ‘hot’ ^3^H-leucine (final concentration 100 nmol leucine L^-1^) for 4 h at room temperature in the dark immediately after sampling. Preliminary experiments show that this was a saturating level of leucine in the offshore water of the SE Mediterranean Sea. After incubation, incorporation was terminated by adding 100 μl trichloroacetic acid (TCA). As a negative control for non-specific binding, another set of triplicates were sampled from a surface layer and treated with TCA immediately after the addition of the radioactive tracer. At the end of each cruise, the samples were processes using the micro-centrifugation protocol and 1ml scintillation cocktail (ULTIMA-GOLD) was added to all samples before counted using TRI-CARB 2100 TR (PACKARD) scintillation counter. A conversion factor of 3 kg C per mole of leucine incorporated and an isotopic dilution of 2.0 were used to calculate the C incorporated (Simon and Azam, 1989).

Net daily photosynthetic carbon fixation rates were estimated using the ^14^C incorporation method (Nielsen, 1952), with several modifications (Hazan et al., 2018). Triplicate 50 ml samples were taken from each depth within the photic zone and from one aphotic depth using sterile vials and kept at surface light and temperature conditions. The ‘dark’ sample served as blank and was kept under the same temperature as the ‘light’ samples. Radioactive spiking was done at ∼08:00 AM the day following of the cruise in order to start a 24h incubation for all samples (including those collected at THEMO-1 station, not shown) at the same time. Early work by Letelier and colleagues (1996) at station HOT showed that prolonged on-deck incubations, similarly to the protocol used in this study, may result in underestimated PP rates as it cannot precisely mimic the temperature and illumination levels in-situ. Our preliminary tests concur with this conclusion and found that ashore incubations underestimate PP rates by up to ∼20% compared to incubations onto a mooring rope tied to the ship (Figure S2). Samples were spiked with 50μl (5 μCi) of NaH^14^CO_3_ tracer and were incubated for 24 h under 3 light regimes: surface illumination (samples from the upper mixing depths), 50% illumination (samples from below the mixing depth to the DCM) and ∼1% illumination (samples from the DCM and below). Shading was performed using neutral density nets, thus changing light intensity but not spectral properties. Water samples were then filtered through GF/F filters (0.7 μm nominal pore size, 25 mm diameter) using low vacuum pressure (< 50 mmHg) and rinsed 3 times with filtered sea water. Filters from each sample were then put in scintillation vials where 50 μl of 32%HCl solution was immediately added in order to remove excess ^14^C-bicarbonate and kept overnight for incubation. After incubation 5 mL scintillation cocktail (ULTIMA-GOLD) was added to the samples and counted using TRI-CARB 2100 TR (PACKARD) scintillation counter. Three random aliquots were counted immediately after the addition of the radiotracer (without incubation) with ethanolamine to serve as added activity measurements.

### 2.3 Picophytoplankton abundance using flow-cytometry

Triplicates water samples (1.5 ml) were collected from each sampling depth, put in cryo-vials (Nunc), and supplemented with 7.5 µl 25% glutaraldehyde (Sigma). Vials were incubated in the dark for 10 min, flash-frozen in liquid nitrogen, and stored in -80 °C freezer. Before analysis, samples were thawed in the dark at room temperature. Each sample was run twice on a BD Canto II flow-cytometer with 2 μm diameter fluorescent beads (Polysciences, Warminster, PA, USA) as a size and fluorescence standard. In the first run three types of phytoplankton cells were identified based on their natural auto-fluorescence: *Prochlorococcus*, *Synechococcus* and picoeukaryotes. Cells were differentiated based on cell chlorophyll (Ex482nm/Em676nm, PerCP channel) and phycoerythrin fluorescence (Ex564nm/Em574nm), and by the size of cell (forward scatter). Before the second FCM, run samples were stained with SYBR Green I (Molecular Probes/ ThermoFisher) to enable counting followed by detection at Ex494nm/Em520nm (FITC channel). This provided counts of the total bacterial population (phytoplankton + heterotrophic bacteria and archaea) as well as a distinction between cells with High or Low DNA content (not shown). Data were processed using FlowJo software. Flow rates were determined several times during each running session by weighing tubes with double-distilled water, and counts of the standard beads were used to verify a consistent flow rate.

### 2.4 Algal pigment markers

Eight litters of seawater were collected from all photic sample depths and one from a dark depth (depth varies between cruises). Water was filtered onto GF/F filters (0.7 μm nominal pore size, 47mm diameter, Waters) using a peristaltic pump until either all 8 L were filtered or the filter became blocked, in which case the volume filtered was recorded. Filters were placed in cryo-vials and flash frozen in liquid nitrogen until they could be stored in a -80 °C freezer. Pigments were extracted in 1ml 100% methanol for 3 h at room temperature and clarified using syringe filters (Acrodisc CR, 13 mm, 0.2 μm PTFE membranes, Pall Life Sciences). Total chlorophyll was measured spectrophotometrically using a NanoDrop 2000c (Thermo Sciences) at 632, 652, 665 and 695 nm, and the concentration of chlorophyll a was calculated (Ritchie 2008). Ultra high-pressure Liquid Chromatography (UPLC) was performed on an ACQUITY UPLC system (Waters) equipped with a photodiode array detector. A C8 column (1.7 μm particle size, 2.1 mm internal diameter, 50 mm column length, ACQUITY UPLC BEH, 186002877) was used. The chromatography method was adapted for UPLC from the LOV method (Hooker et al., 2005). Samples were preheated to 30 °C and column to 50 °C before each run. Running buffers were a 70:30 mixture of methanol and 0.5M ammonium acetate (buffer A) and 100% methanol (buffer B). The program consisted of an isocratic run using a 80:20 mixture of buffers A:B for 2min, followed by a linear gradient to 50:50 for 7 minutes and an increase to 100% solvent B. The flow rate was 0.5ml/min. Pigment standards from DHI (Denmark) were used to identify the UPLC peaks (chlorophyll a, divinyl-chlorophyll a, chlorophyll b, chlorophyll c2, zeaxanthin, beta-carotene, diatoxanthin, fucoxanthin, peridinin, 19’-butanoyloxyfucoxanthin and 19’-hexanoyloxyfucoxanthin). Due to potential degradation of the pigment standards, we present the total chlorophyll measured spectrophotometrically and the pigment ratios within each UPLC run.

## 3 Results

### 3.1 Physical and chemical properties of the water column

Between January 2016 and December 2019, a clear pattern was observed in satellite-derived sea surface temperature at the THEMO-2 location (Figure 1B), which was mirrored during the monthly cruise measurements (February 2018-January 2019, Figure 2A). The measured *in-situ* sea surface temperatures increased by > 11^°^C from the winter minimum of ∼17.9 °C to the summer maximum of 29.1 °C (Figure 2A, Supplementary Figure S3, Table 1). Sea surface salinity also increased from a winter minimum of ∼39.3 psu to a summer maximum of 39.8 psu (Figure 2B, Supplementary Figure S3). Both temperature and salinity minima were higher than the climatological minima (∼15.2 °C and 38.9 psu, measured between 2002-2020 (Herut et al., 2020), suggesting that the sampling period represents a relatively warm and salty year. The temporal changes in sea surface temperature and salinity led to differences in the water density profiles (Figure S1), indicative of stratification of the upper water layer between May and December 2018 (Mixed Layer Depth = 15-49 m) and mixed between February-April 2018 and January 2019 (MLD not determined, see materials and methods) (Table 1). Inorganic nutrient (NO_3_+NO_2_) concentrations began to increase in the mixed layer already during November, suggesting that the stratification had started to erode earlier than observed based on the density profiles, possibly due to short-term mixing events (e.g. (Gunn et al., 2020).

**Figure 2:**
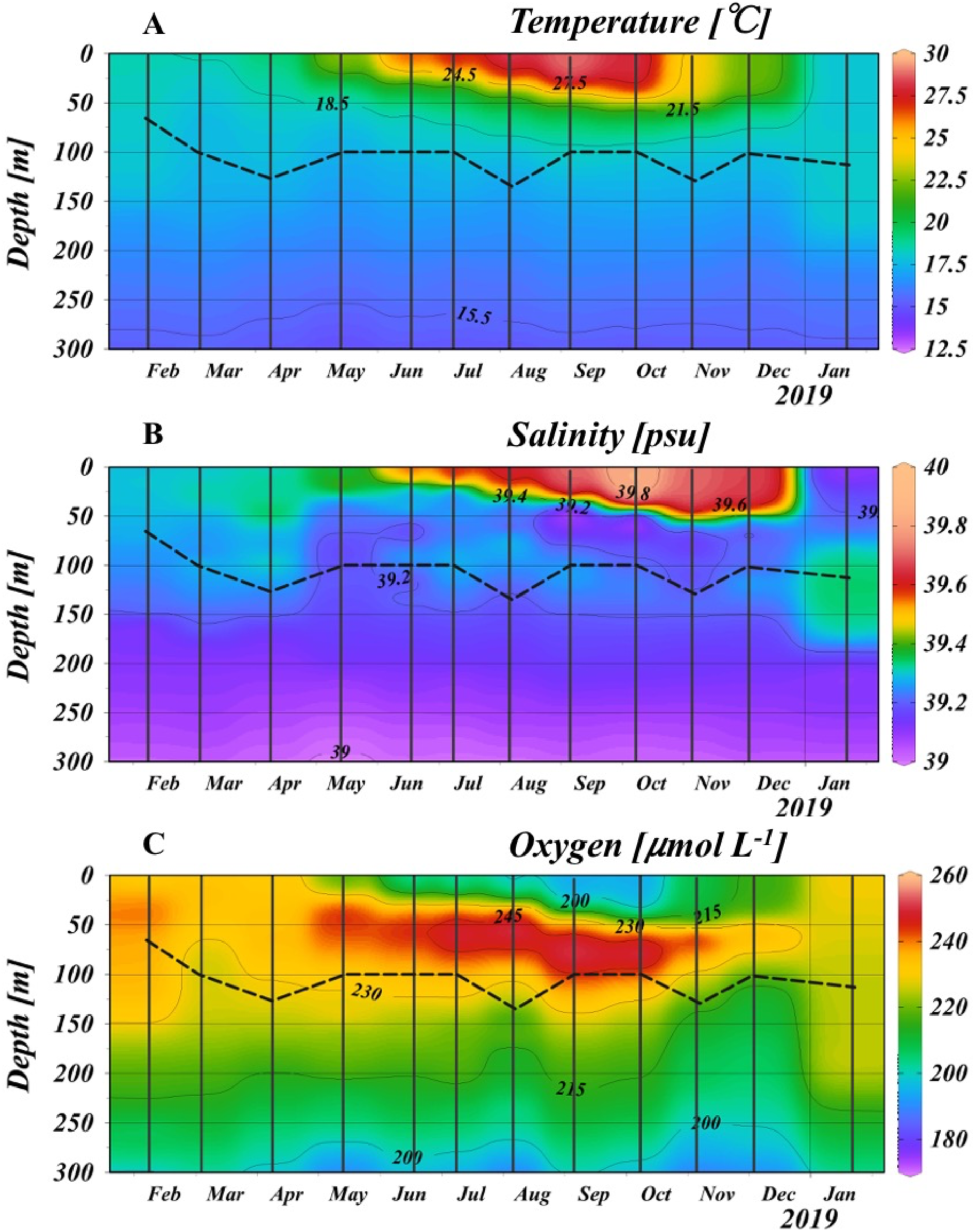
Temperature (A), Salinity (B) and oxygen concentrations (C) at THEMO2 station measured monthly between February 2018 to January 2019. The dashed line represents the depth of the Deep Chlorophyll Maximum (DCM). See Supplementary Figure S3 for the full depth profiles to ∼1,400m.

Dissolved oxygen concentrations were at or above 100% saturation throughout the whole photic layer (0-200 m), ranging from ∼180 µmol L^-1^ to ∼240 µmol L^-1^ (Figure 2C). Where, soluble reactive phosphorus (SRP) concentrations at the surface water were at or close to the limit of detection ∼0.006 μM, (∼6 nM) throughout the year, whereas nitrate+nitrite (NOx) were close to the detection limits from August to November (limit of detection 0.013 μM) but reached 0.3-0.5 μM during the mixed period (Table 1, and see Ben Ezra et al., 2021 for more information).

### 3.2 Chlorophyll a, Primary Productivity and Bacterial Productivity

While satellite-derived sea surface chlorophyll *a* showed a clear seasonal cycle over three years, with lower levels during the stratified period (Figure 1B), the entire photic zone displayed more complex evolution in depth and chlorophyll concentrations (Figure 3A, B). A prominent Deep Chlorophyll Maximum (DCM) was observed year-round, ranging from depths of 60 m (February 2019) to 120 m (April 2019). The concentration of total chlorophyll a at the DCM was always higher than that measured at the surface, and ranged between ∼0.15 *µ*g L^-1^ during February-July, decreasing to ∼0.07 *µ*g L^-1^ during August-November. These values are within the range or somewhat lower than those measured in other studies in the EMS (∼0.09-0.42 *µ*g L^-1^) (Christaki et al., 2001; Yacobi et al., 1995), and are ∼50-80% lower than at BATS (Steinberg et al., 2001). Due primarily to the changes in chlorophyll a at the DCM, the integrated chlorophyll did not follow the same temporal variability as observed with the sea-surface satellites measurements (Figure 1B).

**Figure 3:**
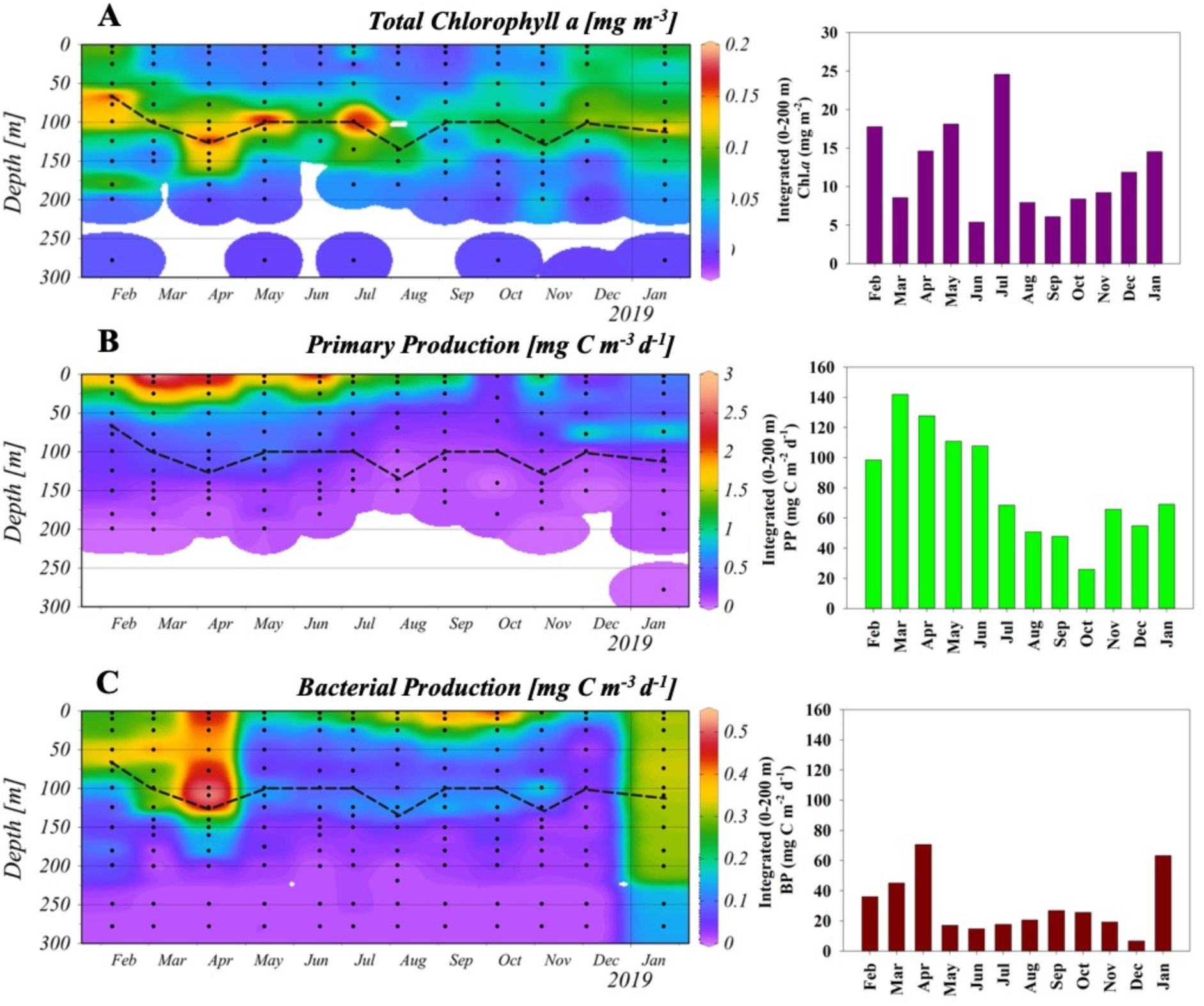
Seasonal changes in depth-resolved (left panels) and depth-integrated (right panels) chlorophyll *a* (A), primary productivity (B) and bacterial productivity (C). Black dots in panels A-C represent sampling points from each cruise. See Supplementary Figure S4 for the full depth profile of BP to ∼1,400m.

Primary productivity (PP) was highest at the surface most of the year (1-3 mg C m^-3^ d^-1^), declining with depth, with no observable maximum during most of the year at the DCM (0.08-0.47 mg C m^-3^ d^-1^, Figure 3B). This is consistent with the chlorophyll maximum in ultra-oligotrophic oceans being decoupled from the PP maximum (Lazzari et al., 2012). An exception to this decoupling was observed during December 2019 and January 2020, where a peak of PP was observed at around 75 m (0.93-0.96 mg C m^-3^ d^-1^), slightly above the DCM (Figure 3B). The observed values at the surface are within the range or somewhat lower than previously measured values in the EMS (typically around ∼2 mg C m^-3^ day^-1^, but with values as high as ∼18 mg C m^-3^ day^-1^ recorded), (Hazan et al., 2018; Rahav et al., 2013). The integrated PP values ranged between 69-142 mg C m^-2^ d^-1^ during the mixing period, and from 48 to 113 mg C m^-2^ d^-1^ during the stratified months (Figure 3B). These values are within the ranges observed in other studies of this region (reviewed in (Berman-Frank and Rahav, 2012; Siokou-Frangou et al., 2010) and discussed below). Our measurements result in an annual PP of ∼32 gC m^-2^, which is lower by ∼50% than most estimates from the EMS (Boldrin et al., 2002; Psarra et al., 2000). These annual estimates highlight the ultra-oligotrophic characteristics of the easternmost Levantine Basin.

During most of the year BP was highest at the surface (∼0.2-0.5 mg C m^-3^ d^-1^, Figure 3C, Supplementary Figure S4), within the range of previous observations at this area (Rahav et al., 2019; Tanaka et al., 2007; Van Wambeke et al., 2000). However, BP differed from PP in several aspects. First, a significant increase in BP was observed during April, with the highest rate observed at ∼110 m where the DCM was detected. Indeed, a smaller secondary peak in BP was observed at a depth corresponding to the DCM throughout the year (Figure 3C).

Second, surface BP increased during summer from a minimum in May to a maximum in September-October, before decreasing again. Third, during January 2019 a large increase was observed in BP, which was more-or-less homogenously distributed throughout the water column down to ∼200 m. Depth-integrated BP ranged from a maximum of ∼70 mg C m^-2^ d^-1^ during April 2018 and January 2019 to a minimum of ∼20 mg C m^-2^ d^-1^ during May-December 2018, within the range of previous measurements ∼10-45 mg C m^-2^ day^-1^ (e.g. (Christaki et al., 2011; Robarts et al., 1996; Van Wambeke et al., 2000).The integrated PP to BP ratio also differed between the mixed and stratified periods. Where, during the mixed season the PP:BP ratio was ∼3:1, while during the stratified period it reached ∼5:1.

### 3.3 Phytoplankton abundance and specific functional groups

Picocyanobacteria (*Prochlorococcus* and *Synechococcus*) were the most abundant picophytoplankton cells throughout the year (Figure 4, Supplementary Figure S5). *Prochlorococcus* was more abundant during the stratified period, and predominated below the mixed layer (Figure 4A). Based on the ratio of divinyll chlorophyll a to total chlorophyll a (Figure 5A), *Prochlorococcus* contributed up to ∼45% of the total phytoplankton biomass at the DCM during late fall (Figure 5A). *Synechococcus* was more abundant during the mixed period and in the surface waters (Figure 4B). Pico-eukaryotes were present at up to 10^4^ cells ml^-1^ during February-June (Figure 4C), but there was no clear group-specific signal in the tested photosynthetic pigments during this period (Figure 5B-D), and thus the specific pico-eukaryotic groups present at this time could not be identified. In contrast, the presence of 19-hexanoyloxyfucoxanthin and 19-butanoyloxyfucoxanthin (19-Hex+19-But), primarily during August to November (Figure 5B), suggests the presence of haptophytes, and this was corroborated with microscopic identification of coccolithophores (see a more detailed discussion in Keuter et al*, in prep*). Finally, peridinin and fucoxanthin peaked during January, suggesting the presence of dinoflagellates and diatoms, respectively (Figure 5C,D).

**Figure 4:**
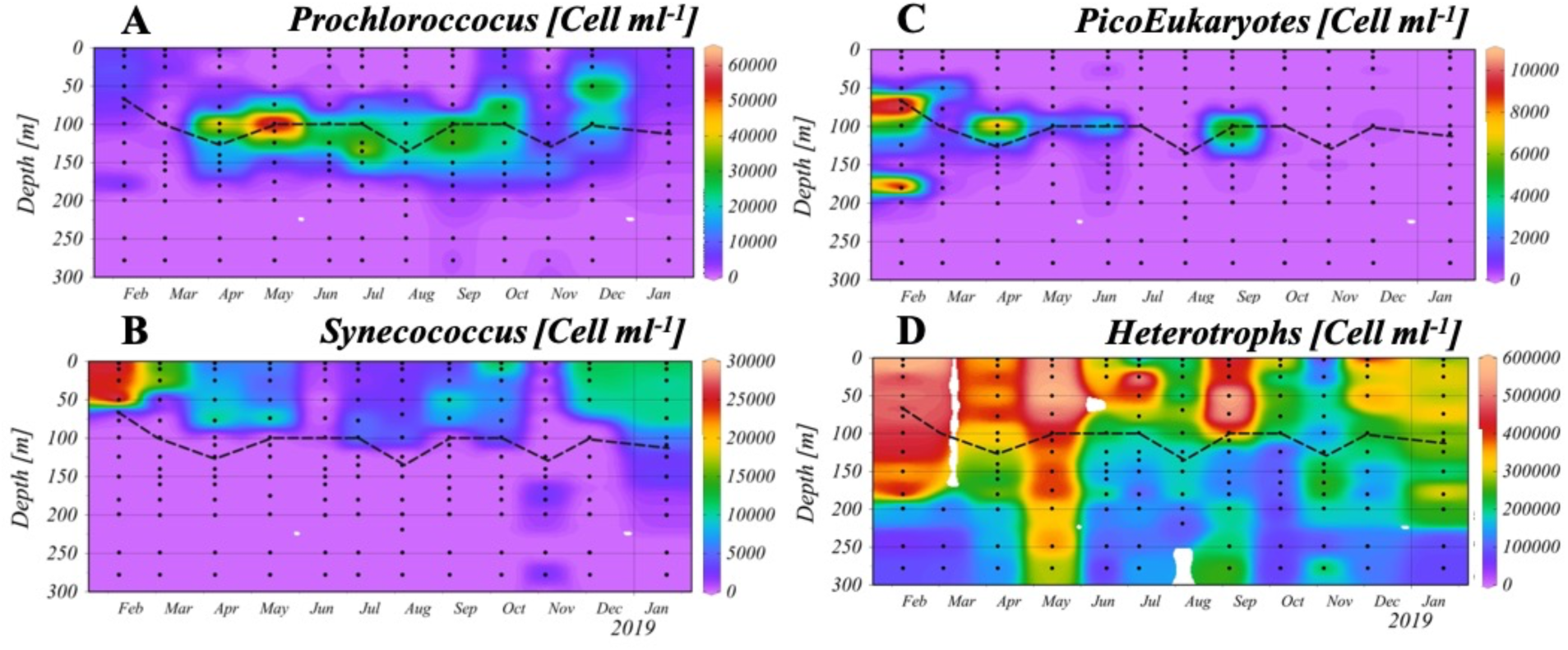
**Annual dynamics of monthly measured picophytoplankton and heterotrophic bacterial abundance:** *Prochlorococcus* (A), *Synechococcus* (B), PicoEukaryotes (C) and heterotrophic bacteria (D) at the THEMO2 station. Black dots represent sample point from each cruise. No data were available from the March cruise for total microbial counts. Note the differences in the color scale for each plot. Dashed line represents the DCM. See Supplementary Figure S5 for the full depth profiles to ∼1,400m.

**Figure 5:**
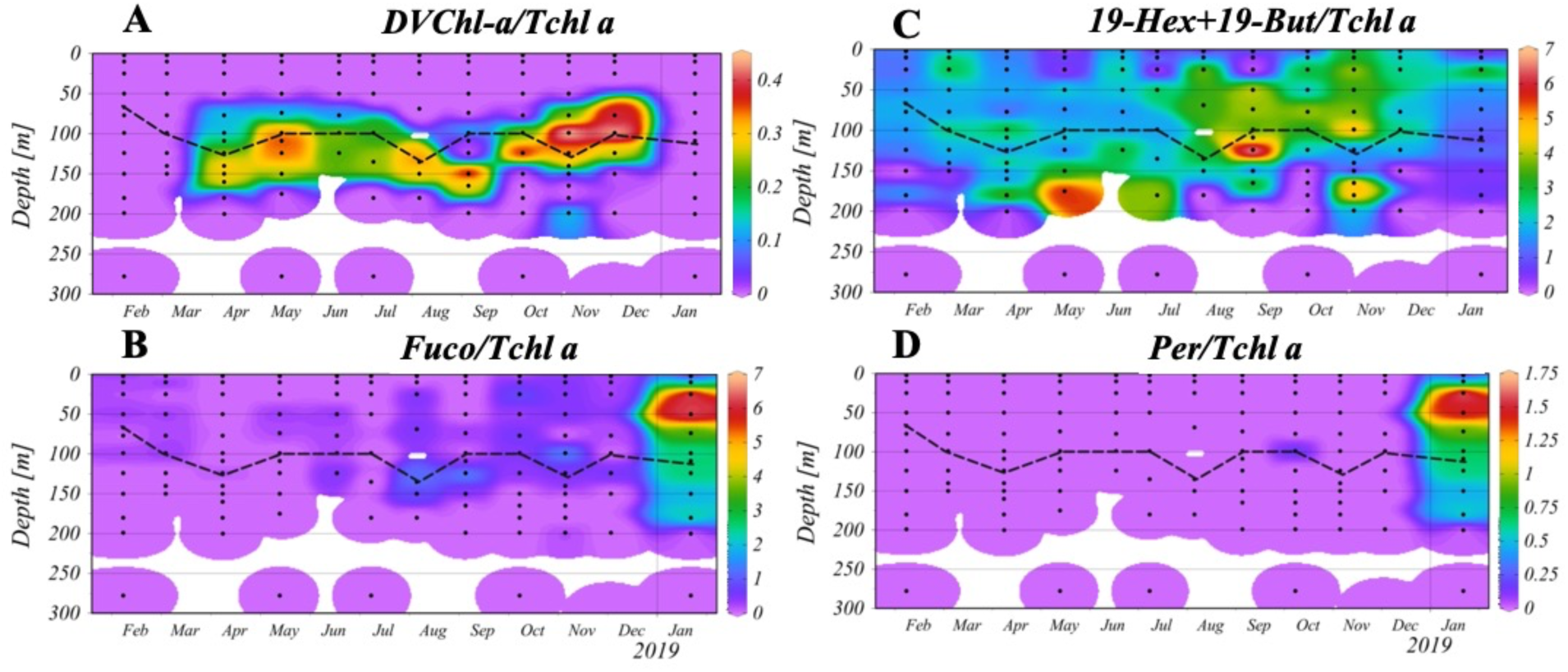
**Monthly changes in the ratios of** major accessory pigments to Total chlorophyll *a*. A) divinyl chlorophyll *a* (DVChl-a). B) Combined 19-hexanoyloxyfucoxanthin and 19-butanoyloxyfucoxanthin (19-Hex+19-But). C) Fucoxanthin (Fuco). D) Peridinin. Black dots represent sampling depths. Dashed line represents the DCM.

Throughout the year, total prokaryotic microbial counts (Sybr-stained cells, thus including cyanobacteria, heterotrophic bacteria and archaea) ranged between 3 ×10^5^ to 6×10^5^ cells ml^-1^, with higher values observed during February and May (Figure 3SB). Heterotrophic bacteria were much more abundant than the combined phototrophs, the latter forming 2.4-8.5% of the total Sybr-stained cell counts (Figure 4D). Below the photic layer, cell counts were typically lower, ranging from 1.9×10^4^-4.5×10^5^ cells ml^-1^ (Supplementary Figure S5).

## 4 Discussion

### 4.1 How oligotrophic is the EMS compared to other oligotrophic oceans?

The EMS has previously been suggested to be one of the most oligotrophic marine systems on Earth, including a claim for the “deepest Secchi-depth world record” - 53 m during summertime (Berman et al., 1984). A comparison of published values of PP, including those measured in this study, supports this notion, with the median integrated PP being ∼66% lower than Bermuda, the Red Sea and the Western Mediterranean Sea (WMS), and ∼80% lower than station ALOHA (HOT) (Figure 6A). Our estimates of PP during the SoMMoS cruises are among the lowest in the EMS (Figure 6A), yielding an annual PP of ∼32-39 gC m^-2^ y^-1^, although similar values have previously been reported (Dugdale and Wilkerson, 1988). The annual PP during the SoMMoS cruises is approximately half of that reported above the continental slope of the Cretan Sea, ∼59 gC m^-2^ y^-1^, (Psarra et al., 2000), as well as in other more western locations in the EMS, ∼62 gC m^-2^ y^-1^ (Boldrin et al., 2002). It should be noted, though, that our incubation approach do not precisely mimic the *in-situ* illumination and temperature at the time of sampling, which, based on our previous work, may underestimate the actual PP rates by up to ∼20% (Figure S2). Under these circumstances, the annual PP we report may reach ∼39 gC m^-2^ y^-1^, approximately ∼33% lower than previously reported from the EMS (Psarra et al., 2000; Boldrin et al., 2002). Moreover, comparison between different studies that used similar, yet not identical, methodology for PP estimates, should be done with care. For example, in the comparison presented in Figure 6A many of the studies measured PP from dawn to dusk while in this study we used longer incubations which account for the whole day (net PP). These differences may also, partly, account for the changes between the different oceanic locations, yet even with these caveats taken into account the PP in the EMS is still lower than other oceanic regions (Figure 6A).

**Figure 6:**
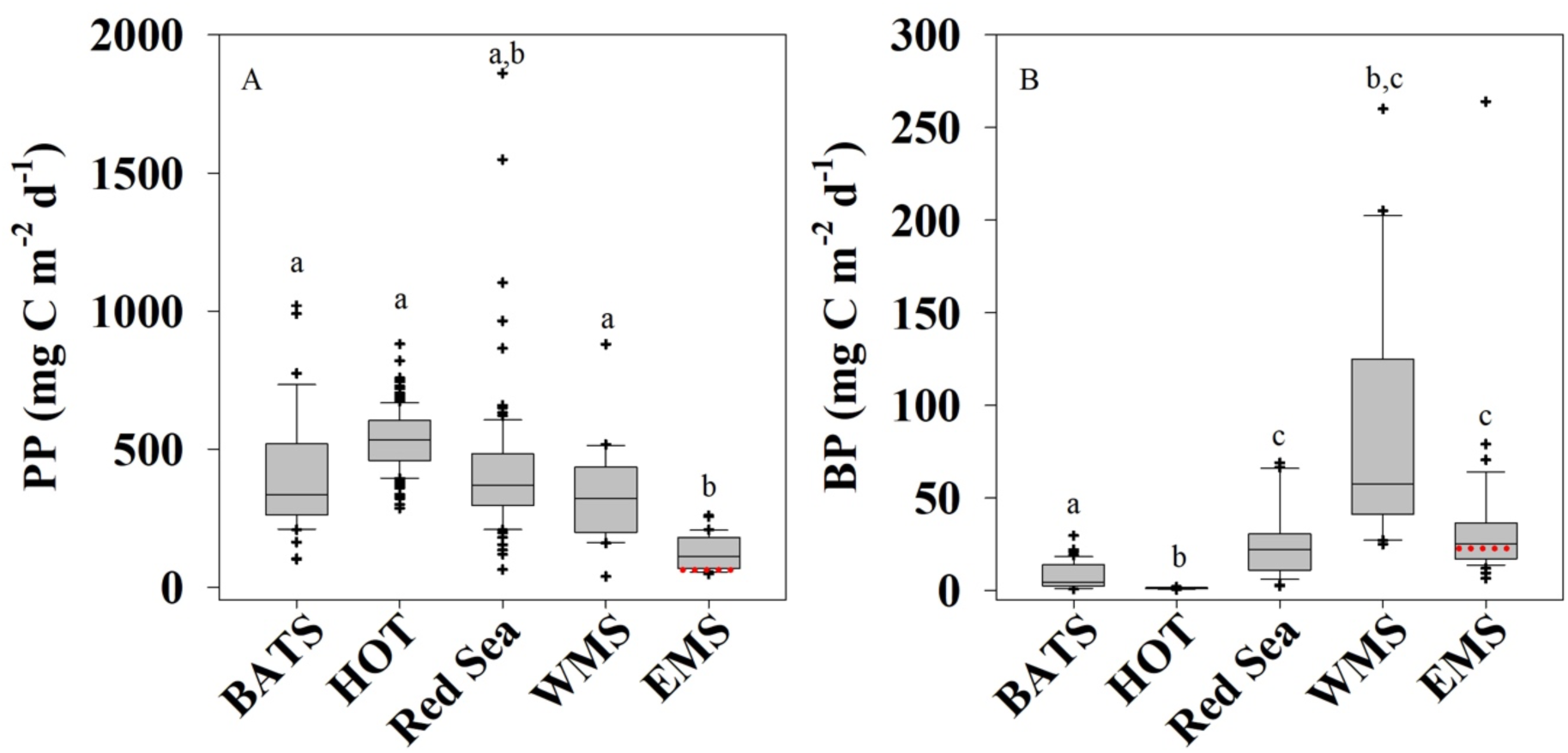
**Comparative literature analysis of the integrated primary and bacterial productivity** (PP and BP, respectively) in the EMS and other well-studied oligotrophic locations (Bianchi et al., 1999; Bonnet et al., 2011; Casotti et al., 2003; Christaki et al., 2011; Decembrini et al., 2009; Fernández et al., 1994; Gasol et al., 1998; Gaudy et al., 2003; Hazan et al., 2018; Ignatiades et al., 2002; Lemée et al., 2002; Lohrenz et al., 1988; Morán and Estrada, 2001; Moutin and Raimbault, 2002; Pedrós-Alió et al., 1999; Rahav et al., 2019; Eyal Rahav et al., 2013; Robarts et al., 1996; Siokou-Frangou et al., 2002; Van Wambeke et al., 2004; Vidussi et al., 2000; Wambeke et al., 2002; Zervoudaki et al., 2007). Box-Whisker plots show the interquartile range (25th–75th percentile) of the data set. The horizontal line within the box represents the median value. The red dashed line in the EMS boxes shows the median values for the SoMMoS cruises described here. The letters above the box-plots represent significant differences (ANOVA, p < 0.05) for mean values between sampling sites. The BP compilation includes only measurements obtained using the ^3^H-leucine incorporation method. Note the different Y axis.

The values of integrated chlorophyll (Table 1, Figure 2A) and phytoplankton cell counts are also somewhat lower than previously-published measurements from the same region (Figure 4, Figure 7) (Berman et al., 1984; Robarts et al., 1996). Both the temperature and salinity minima during the SoMMoS cruise series were higher than the 2002-2020 climatological average (Herut et al., 2020), and thus we currently cannot determine whether the lower PP values measured during this annual cruise series are due to inter-annual variability, or whether these are due to the possible underestimation of the PP values discussed above (section 2.2). Nevertheless, it is clear that the EMS is one of the most oligotrophic locations on Earth, as defined using primary production, and that key seasonal dynamics are similar between studies performed a decade apart (compare Figures 7D, E).

**Figure 7:**
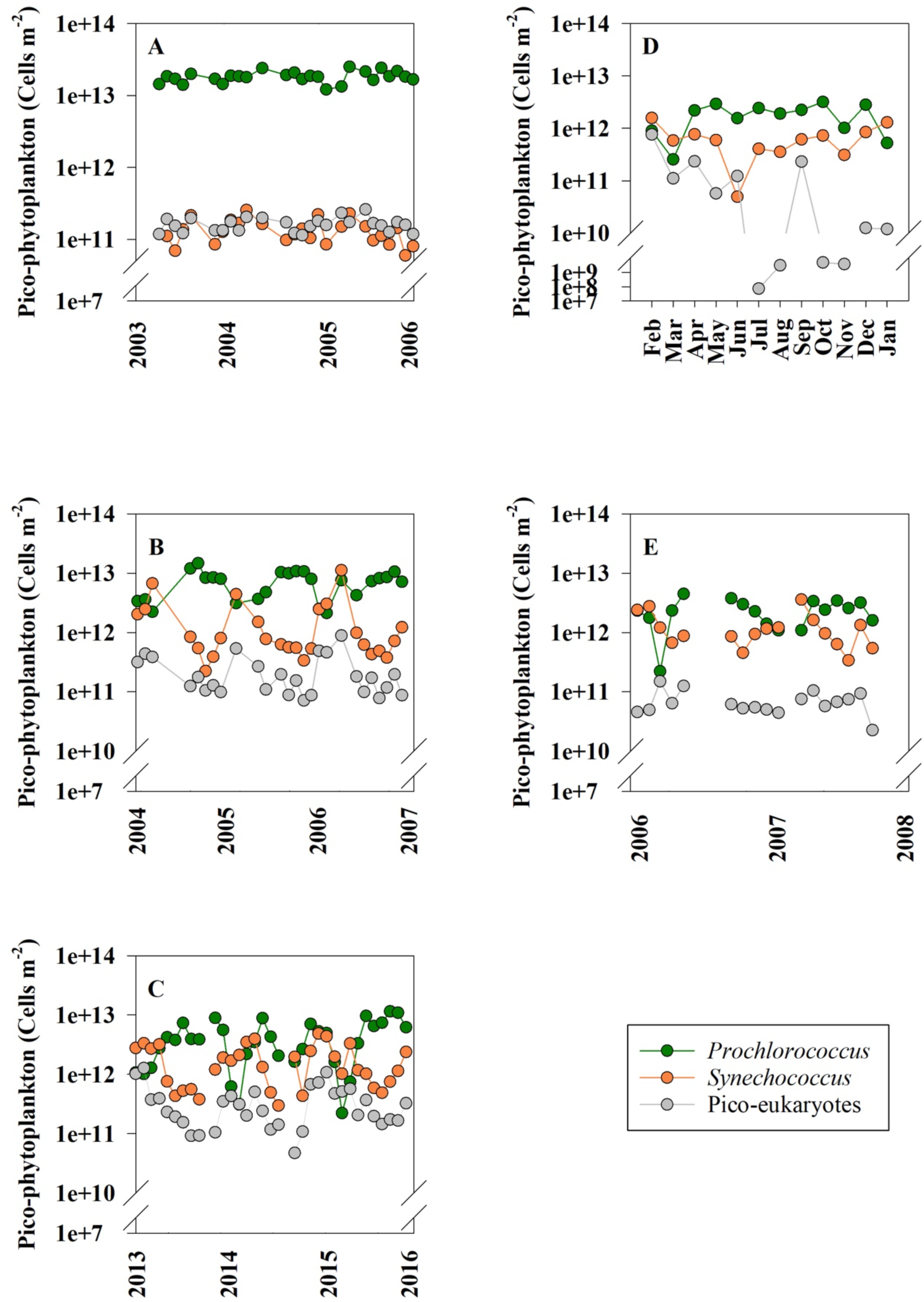
Integrated cell count time series of *Prochlorococcus* (green), *Synechococcus* (orange) and picoeukaryotes (gray) at stations HOT (A), BATS (B), the Northern Red Sea (C), and the EMS (D, E – this study and (Yogev et al., 2011) respectively). Abundance measurements were carried out using flow cytometry and are presented in a log scale. Data for stations HOT and BATS were compiled from (Malmstrom et al., 2010) and from the Northern Red Sea from the Eilat National Monitoring Program (Shaked and Genin, 2017).

In addition to low PP, the EMS exhibits low sinking fluxes of particulate organic carbon (POC). The POC flux measured from the bottom of the photic layer (180 m) at the DeepLev mooring station, which is in the same region as the THEMO-2 site (Figure 1A, ∼10Km north of our study site, Lat: 32.820 N; 34.380 E), is ∼0.2 gC m^-2^ during the summer months (May-September 2018) and ∼0.8 gC m^-2^ per year (Alkalay et al., 2020). This POC flux to the aphotic layer is less than 2% of the PP measured during summertime (∼12 gC m^-2^ in 5 months) or 2.5% from the annual PP rates (∼32 gC m^-2^ y^-1^). The fraction of PP exported as particulate matter in the EMS is therefore lower than that reported in other oligotrophic systems; i.e., HOT and BATS stations ∼5% (Karl and Church, 2014; Steinberg et al., 2001). This low contribution provides an additional biogeochemical implication of the extreme oligotrophic conditions of the EMS. It also points to that the dissolved organic carbon fraction derived from PP is rapidly recycled in the photic layer by heterotrophic/mixotrophic bacteria as often observed in LNLC environments, as reviewed in (Santinelli, 2015). We note, however, that estimates of export production based on nitrate+nitrate loss from the photic zone, calculated from the same data, are significantly higher (172 mmol N m^-2^ year^-1^, corresponding to ∼13.6g C m^-2^ year^-1^, presented in a companion paper, Ben Ezra et al., 2021). The reason for this discrepancy is unclear, but may be due to over-estimation of the export production rates (e.g. if part of the nitrate+nitrite was taken up by heterotrophic organisms) and/or underestimation of the sinking carbon flux from sediment traps, and/or non-Redfield C:N conversion factors that may prevail in the EMS. Moreover, such an underestimation may be due to sinking flux from organisms too large to be caught in the sediment traps such as fish or jellyfish (Edelist et al., 2020).

In contrast to PP, integrated BP in the EMS was higher than estimates at BATS and HOT, and similar to the Red Sea (Figure 6B, which compares only measurements obtained using the ^3^H-leucine incorporation method). Yet, values obtained in the South Pacific Gyre, 43-61 mg C m^-2^ d^-1^, (Berman-Frank et al., 2016) do come close to our measurements in the EMS. Thus, differences in PP between oligotrophic regions are not necessarily coupled to BP measurements. Previous studies, including from HOT and BATS, have shown that BP and PP are decoupled across multiple time-scales (e.g hours-months) (Viviani and Church, 2017). Similarly, PP and BP were uncoupled in the cyclonic Rhodes Gyre and the anti-cyclonic Cyprus Eddy of the EMS (Rahav et al., 2013) and also in our study when BP increased in surface waters between June and November, while PP rates declined (Figure 3). One potential reason for this discrepancy is that PP is often measured on filtered samples (particulate PP), and thus does not include the fraction of dissolved organic carbon (DOC) fixed through PP that is released from the cells due to exudation or lysis (Viviani et al., 2015).

When phytoplankton have sufficient carbon and light for photosynthesis yet nutrients such as P and N are limiting, photosynthesis is uncoupled from growth and high concentrations of DOC are often released to the environment (Berman-Frank and Dubinsky, 1999). Previous studies from natural samples and from cultures of the numerically-dominant phytoplankton in the EMS, *Prochlorococcus* and *Synechococcus*, suggest a potentially high but variable fraction of released DOC (typically 20-70% of PP, with one study suggesting >90% in lab cultures, (Roth-Rosenberg et al., 2020; Viviani et al., 2015). It is currently unclear whether autochthonous production (that is, local PP) is sufficient to support the growth requirements of heterotrophic bacteria, given what is known about the growth efficiency of bacteria. A study from the Cretan Sea suggesting that this is possible if the release of DOC from phytoplankton exceeds 40% of PP (Anderson and Turley, 2003).

The EMS is affected by coastal intrusions of relatively chlorophyll-rich waters (Efrati et al., 2013), which could contribute also additional DOC, yet the importance of these processes in determining BP and PP requires additional studies, as they occur on temporal and spatial scales not captured by the SoMMoS cruise series. Such studies are also needed to understand the relative contribution of dissolved compared to particulate carbon to export processes in oligotrophic regions.

An additional explanation for the uncoupling between PP and BP was previously suggested following an *in-situ* phosphorus addition experiment at an anti-cyclonic eddy in the EMS, where orthophosphate addition led to chlorophyll *a* decrease while heterotrophic biomass and activity increased (Thingstad et al., 2005). The authors postulated two possible scenarios to explain their observation; 1) fast growing bacteria out-competed phytoplankton for the added phosphorus. Then, the accumulated heterotrophic biomass was quickly channeled toward larger consumers (the ‘bypass hypothesis’). 2) luxury uptake of P, mainly by heterotrophic bacteria (and to a lesser extent by small-size picophytoplankton), formed a phosphorus-rich diet for grazers, resulting in increased egg production (the ‘tunneling hypothesis’).

Additionally, a scenario where an increase in temperature drives increased growth by heterotrophic bacteria (Luna et al., 2012), resulting in a draw-down of inorganic nutrients, could explain the increased BP in surface waters during summer, when no increase in PP was observed (Figure 3).

### 4.2 Who are the main phytoplankton in the EMS?

During the thermally mixed period, *Synechococcus* were more abundant than *Prochlorococcus* at the upper ∼100 m (Figures 4). This period was characterized by measurable N at the surface while P concentrations were still extremely low – possibly causing P limitation (December-July, Table 1, Ben Ezra et al., 2021). In contrast, the extreme surface nutrient scarcity during the stratified period, when both N and P were below the limit of detection and were potentially co-limiting (Ben Ezra et al., 2021), likely contributed to the dominance of the small-size cyanobacterium *Prochlorococcus* over the somewhat larger-cell *Synechococcus* (Figures 4). It is generally accepted that, as the nutrient concentrations decrease (i.e., the water becomes more oligotrophic), so does the average size of phytoplankton (Margalef and Kinne, 1997). Additionally, while most *Synechococcus* genomes encode the genes required for nitrate uptake and utilization, this trait is found only in a subset of the *Prochlroococcus* genomes (Berube et al., 2018), and single-cell analysis suggests that more *Synechococcus* cells in nature utilize nitrate compared to *Prochlorococcus* (Berthelot et al., 2019). Thus, the type of nutrient limitation, P or N+P, (Krom et al., 1991; Thingstad et al., 2005; Zohary et al., 2005) may also affect the temporal dynamics of these two clades in the water column.

The abundances of these two cyanobacterial groups also differed with depth. *Synechococcus* was more abundant in the surface waters than *Prochlorococcus* (Fig 4). This was especially evident from May to September when *Prochlorococcus* cells could not be identified in the surface layer using either flow cytometry or pigment analysis (Figures 4 and 5). While the low *Prochlorococcus* abundance at the surface could be partly due to the limit of detection of the flow cytometer used, it is supported by the lack of observed DVChl-a (Figure 5a) and by previous genetic analyses (Rosenberg et al., 2020). It has been suggested that *Prochlorococcus’* unique photosynthetic pigments (DVChl-a and DVChl-b) provide a competitive advantage for these organisms at deeper layers of the water column, leading to a niche separation between these two pico-cyanobacteria (Moore et al., 1995). The observed niche (depth) separation between *Prochlorococcus* and *Synechococcus* has been previously observed including in the first description of *Prochlorococcus* from the North Atlantic and North Pacific (Chisholm et al., 1988), and the Eastern South Pacific (BIOSOPE cruise, (Huang et al., 2015)). However, this is not a universal observation. At the HOT station in the North Pacific, *Prochlorococcus* and *Synechococcus* often show similar depth distributions, both being more abundant at the surface, with *Prochlorococcus* more abundant numerically and in terms of biomass (van den Engh et al., 2017). Similarly, in the Indian Ocean during May-June 2003 *Prochlorococcus* dominated the surface waters whereas *Synechococcus* were much less abundant (Huang et al., 2015). Indeed, a global analysis of >35,000 quantitative measurements of *Prochlorococcus* and *Synechococcus* found that *Prochlorococcus* were often more abundant than *Synechococcus* at high PAR and temperature values representative of surface waters in tropical waters (Flombaum et al., 2013). Thus, the reasons why *Prochlorococcus* and *Synechococcus* partitioned by both depth and season in the EMS, and why this is not necessarily observed elsewhere, remain unclear. Other factors probably also impact distribution, as was suggested for surface populations of *Prochlorococcus* that appeared to be negatively affected by atmospheric deposition through biological (e.g., airborne viruses or other biological agents that infect the cells) and chemical (e.g., trace-metals toxicity) processes (Rahav et al., 2020).

While the pico-cyanobacteria *Prochlorococcus* and *Synechococcus* were the most numerically abundant phytoplankton at THEMO-2, when biomass is examined the contribution of picoeukaryotes can be as high as that of the two pico-cyanobacteria, or even higher (Supplementary Figure S6, and see below). A previous study of the diversity of pico-phytoplankton in the EMS showed that pico-eukaryotes were not very common in terms of their DNA sequences or flow cytometry signals, but were often dominant in terms of RNA sequences, one potential proxy for photosynthetic activity (Man-Aharonovich et al., 2010). In that study, pico-eukaryotes were suggested to comprise up to 60% of the photosynthetic picoplankton biomass in surface waters, and were mainly composed of Haptophytes (primarily *Prymnesiophytes*) and *Stramenopiles* (Man-Aharonovich et al., 2010). In agreement with these observations, the photosynthetic pigment 19’-hexanoyloxyfucoxanthin (19’-hex), which is considered (together with 19’-butanoyloxyfucoxanthin, or 19’-but) to be a diagnostic pigment for *prymnesiophytes*, was found year-round in the water column, but was more abundant relative to the total phytoplankton biomass (i.e. in relation to total chlorophyll) during August-October (stratified period). Thus, both molecular evidence (Man-Aharonovich et al., 2010) and biochemical evidence (pigment analysis) point to the importance of prymnesiophytes in the EMS. A companion study from the SoMMoS cruises provides an overview of seasonal dynamic and impact of calcified haptophytes (coccolithophores) during this yearly survey (Keuter et al, *in prep*).

Interestingly, we often observed 19’-hex around or below the DCM, including at depths where light intensities are very low and photosynthesis may not provide enough carbon or energy to support growth (Fig 5). Recent studies have unveiled that haptophytes, including coccolithophores, are mixotrophic - capable of acquiring prey by phagotrophy or organic compounds by osmotrophy in addition to photosynthesis (e.g. (Avrahami and Frada, 2020; Godrijan et al., 2020; Tillmann, 1998). Indeed, haptophytes can contribute to large fractions of total bacteriovory in marine settings (Frias-Lopez et al., 2009; Unrein et al., 2014). Further analysis of the DNA samples collected on the SoMMoS cruises may help identify mixotrophy within the phytoplankton clades found at the base of the photic layer (e.g. (Yelton et al., 2016) and help illuminate the metabolic capacities required to support phytoplankton growth in this environment.

Pigments associated with diatoms and dinoflagellates, which are often dominant phytoplankton groups (Andersen et al., 1996), were observed at relatively high concentrations only during January 2019 (Figure 5C, D). At this time DVChl-*a* measurement were below the detection limit in the water column, suggesting a succession from pico-cyanobacteria to larger eukaryotic organisms, which occurred relatively rapidly - over several weeks. This occurred concomitantly with a drop in surface temperature (Figure 2A) and thus a break in stratification, an increase in nitrate+nitrite (Ben Ezra et al., 2021), and a shift in coccolithophore population towards r-selected species such as *Emiliania huxleyi* which are indicative for a higher nutrient regime (Keuter et al*, in prep*). While chlorophyll-*a* concentrations increased both at the surface (including by satellite sensing, Figure 1B) and at the DCM (Figure 3A), no concomitant increase was measured in PP (Figure 3B). Yet, a major increase was observed in BP (Figure 3C). As described in detail in the materials and methods, our measurements of PP likely underestimate productivity at depth, and the pigments were observed primarily below the mixed layer. It is possible that the phytoplankton bloom actually occurred several days before our cruise, and that the pigments observed are from stressed or dead phytoplankton. In this case, the major increase in BP observed at the time may represent heterotrophic bacteria utilizing dissolved and particulate organic carbon produced during the bloom.

### 4.3 Phytoplankton composition and seasonality in the EMS in comparison to HOT, BATS and the Red Sea

The north Atlantic and Pacific gyres, the northern Red Sea and the EMS are all considered Low-Nutrient Low-Chlorophyll marine systems, with phytoplankton numerically dominated by pico-cyanobacteria throughout most of the year. Nevertheless, there are differences in the compositions and the seasonal dynamics of the different phytoplankton clades among these LNLC location (Fig. 7, Sup Fig. 5). Specifically, BATS, the Red Sea and the EMS all show seasonal succession patterns in the abundance of the main phytoplankton groups, although there are subtle differences between these sites: i) The period when integrated *Synechococcus* numbers are higher than *Prochlorococcus* is consistently longer in the Red Sea and EMS (3-4 months) compared to BATS (1-2 months); ii) *Prochlorococcus* reached higher absolute cell abundances during the stratified period at BATS (∼10^13^ cells m^-2^) compared to the Northern Red Sea or the EMS (∼2.5×10^11^ cells m^-2^); iii) In terms of calculated biomass, the Red Sea is dominated by pico-eukaryotes for much of the year, whereas in BATS and the EMS the contribution of *Prochlorococcus*, *Synechococcus* and pico-eukaryotes to the total biomass is similar, and the dominant groups change over the year (Supporting Fig S6).

The absolute nutrient concentrations and the identity of the limiting nutrient(s) are different between these locations. For example, BATS is typically considered P limited, whereas the EMS is potentially P limited during winter and co-limited by N and P during summer (Zohary et al., 2005). There may also be differences in the length of the mixed period, its intensity, or the composition of the mixed water (e.g. deep water in the EMS have an N:P ratio of ∼28:1, compared to the canonical 16:1 Redfield ratio (Redfield, 1934). Future work may use the observed differences in phytoplankton dynamics as a sensitive “readout” of the system, enabling a better understanding of the oceanographic processes underlying the differences between BATS, the Red Sea and the EMS.

In contrast to BATS, the Red Sea and the EMS, seasonality at HOT in is much less pronounced (Figure 7). *Prochlorococcus* are about 2-orders of magnitude more abundant than *Synechococcus* and pico-eukaryotes, and are also the most dominant in terms of biomass (Sup Fig. 5). The drivers of phytoplankton temporal dynamics at HOT are thus likely fundamentally different compared to BATS, the Red Sea and the EMS. It has been suggested that a-periodic events such as mesoscale eddies, Fe deposition from winds, and biological processes such as dinitrogen fixation, may be major drivers of phytoplankton dynamics at HOT (Karl and Church, 2014) and less so in the EMS (Berman-Frank and Rahav, 2012; Yogev et al., 2011).

## 5 Closing remarks

The year of data presented here show that the EMS is one of the most oligotrophic regions in the global oceans, as estimated based on PP (32 mg C m^-2^ year ^-1^), chlorophyll a (median ∼10 mg m^-2^), and dominance of small-size picophytoplankton with overall low abundances compared to other LNLC regions (in some cases ∼2 orders of magnitude lower). In contrast, BP in the EMS is higher than at HOT and BATS, and similar to the Red and Western Mediterranean seas. This suggests that the system is overall heterotrophic, and that different factors might limit phytoplankton and heterotrophic bacteria (e.g. (Thingstad et al., 2005).

Despite these observations, there is a clear seasonal cycle in the phytoplankton populations and their activities, reminiscent of BATS and the Northern Red Sea (but not of HOT), suggesting that mixing (and potentially other seasonal processes such as dust deposition) drive similar processes across many, but not all, LNLC regions.

Our results also highlight important knowledge gaps regarding the EMS. For example, we do not know to what extent the bacterial populations or their physiological traits (e.g. bacterial carbon demand or growth efficiency) differ seasonally or vertically in the EMS, as well as compared to other LNLC locations. Detailed studies are also lacking of higher trophic levels in this region (dominant micro-grazer species, diversity, biomass, grazing rates, etc.). These are critical for understanding ecosystem dynamics including top-down vs. bottom-up regulations etc. Biogeochemical characterization of the EMS will also benefit from more explicit characterization of carbon-per-cell estimates, used to derive the biomass contributions from flow-cytometry counts (i.e. Supplementary Figure S6) can vary depending on taxonomy and physiological state (Kirchman, 2013). Future studies of these aspects, together with other data emerging from the SoMMoS cruise series and the ocean observatories in the EMS, should provide the needed background for governments and other stakeholders to employ science-based environmental policies in this rapidly changing region.

## Acknowledgements

We thank the captains and crew of the R/V Mediterranean Explorer (EcoOcean) and R/V Bat-Galim (IOLR), Oshra Yosef, Elad Rachmilovitz, Guy Sisma-Ventura for help with the sampling. The SoMMoS ship-time was funded by the Leon H. Charney School of Marine Sciences with help from EcoOcean and IOLR. This study was supported by grant RGP0020/2016 from the Human Frontiers Science Program (to DS), by grant number 1635070/2016532 from the NSF-BSF program in Oceanography (NSFOCE-BSF, to DS), and partly by the Israel Science Foundation (grants number 1211/17 to ER and BH and 996/08 to I. B-F and B.H). This study is in partial fulfilment of the M.Sc. thesis of Tom Reich (University of Haifa).

## Author Contributions

MDK, YL, ER and DS initiated and designed study; TR, TBE, NB, DRR, OB, MDK, ER and DS collected samples; TR, TBE, NB and AT analyzed samples with help from MDK, DA, DRR and SG; TR, ER and DS wrote the manuscript with input from all co-authors.

