## Supplementary material for "A year in the life of the Eastern Mediterranean: Monthly dynamics of phytoplankton and bacterioplankton in an ultra-oligotrophic sea": Supplemetary Info

### Supplementary Information

#### Table S1 : *THEMO_ALL.xlsx* (supplied as digital file)

File contains: Cruise meta data, Abiotic parameters, Cell counts, PP, BP, Pigment data.

#### Figure S1 : Temperature, Salinity and Density depth profiles (to 300m) for each of the 12 cruises to the THEMO-2 station.

#### Figure S2: Physical parameters (temperature salinity and density), across the entire water column at THEMO-2 (down to ~1,500m).

#### Figure S3: Cell counts of phytoplankton groups and of total cells across the entire water column at THEMO-2 (down to ~1,500m)

**Figure S4**: Profiles of bacterial production across the entire water column at THEMO-2 (~1,500m)

**Figure S5**: Integrated biomass time series of the calculated biomass of *Prochlorococcus* (green), *Synechococcus* (orange) and picoeukaryotes (gray) at stations HOT (A), BATS (B), the Northern Red Sea (C), and the EMS (D – this study and E - Yogev et al., 2011). Biomass measurements were calculated from the flow cytometry counts and assuming a *Prochlorococcus* biomass of 53 fgC cell^-1^, *Synechococcus* biomass of 175 fg C cell^-1^ and pico-eukaryotes biomass of 2100 fgC cell^-1^ (Campbell 2001). Data for stations HOT and BATS were compiled from (Malmstrom et al. 2010) and from the Northern Red Sea from the Eilat National Monitoring Program 2017.

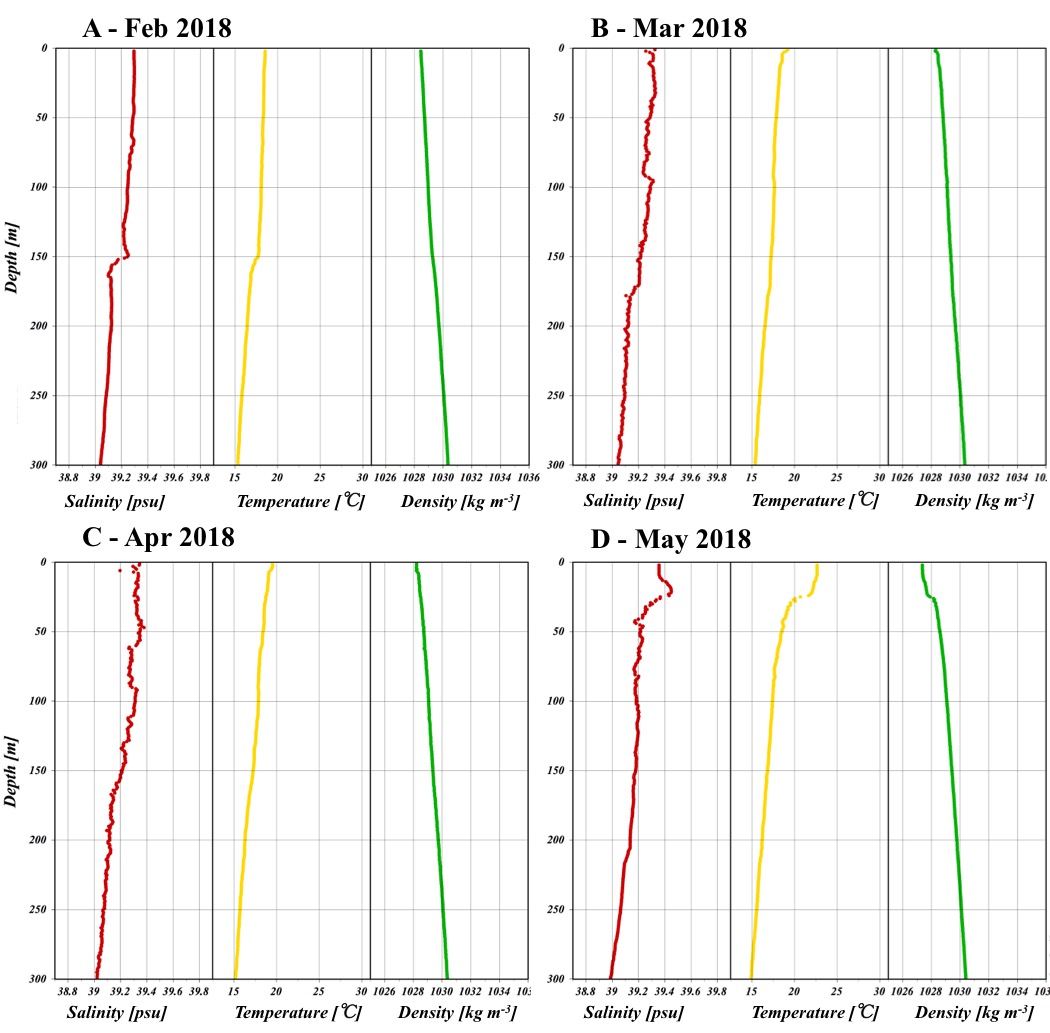

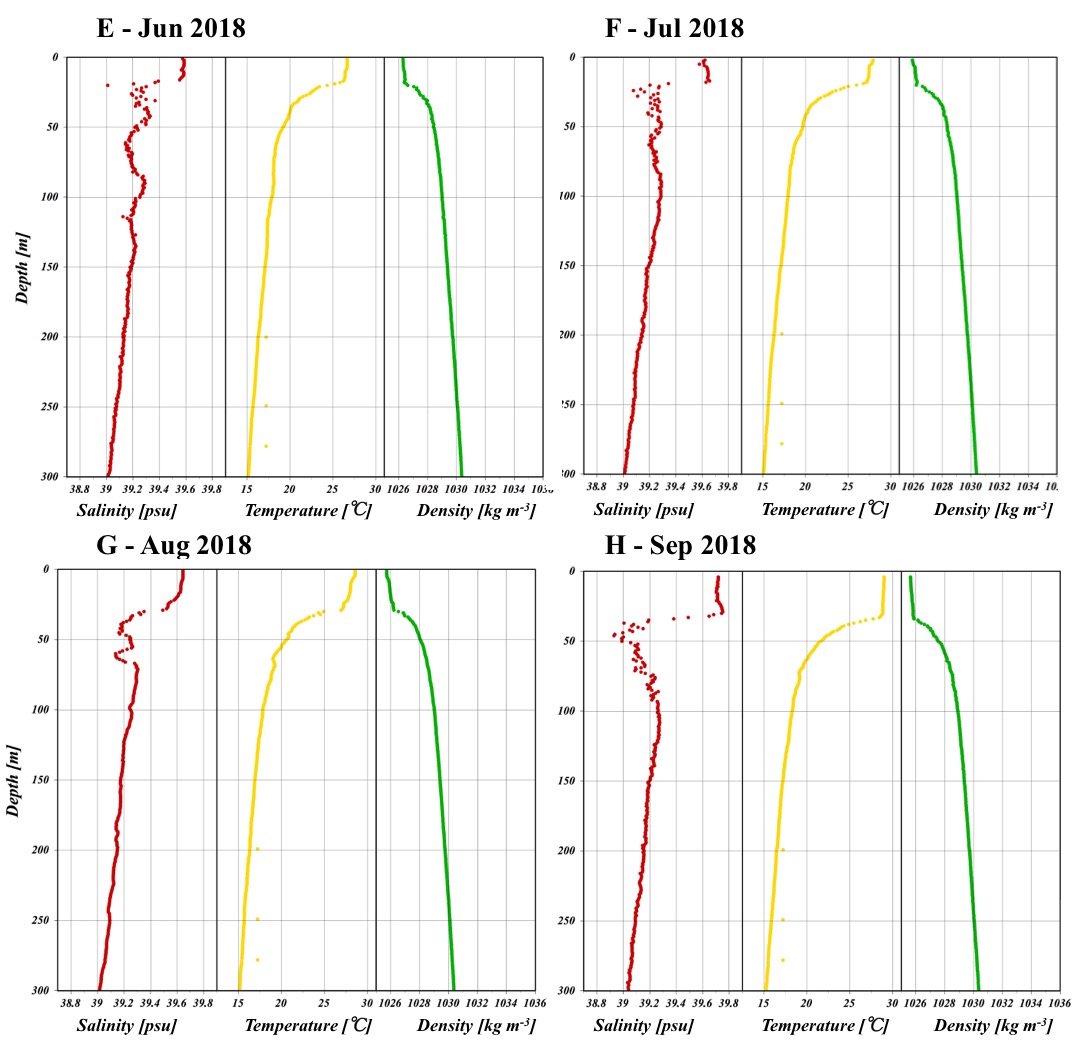

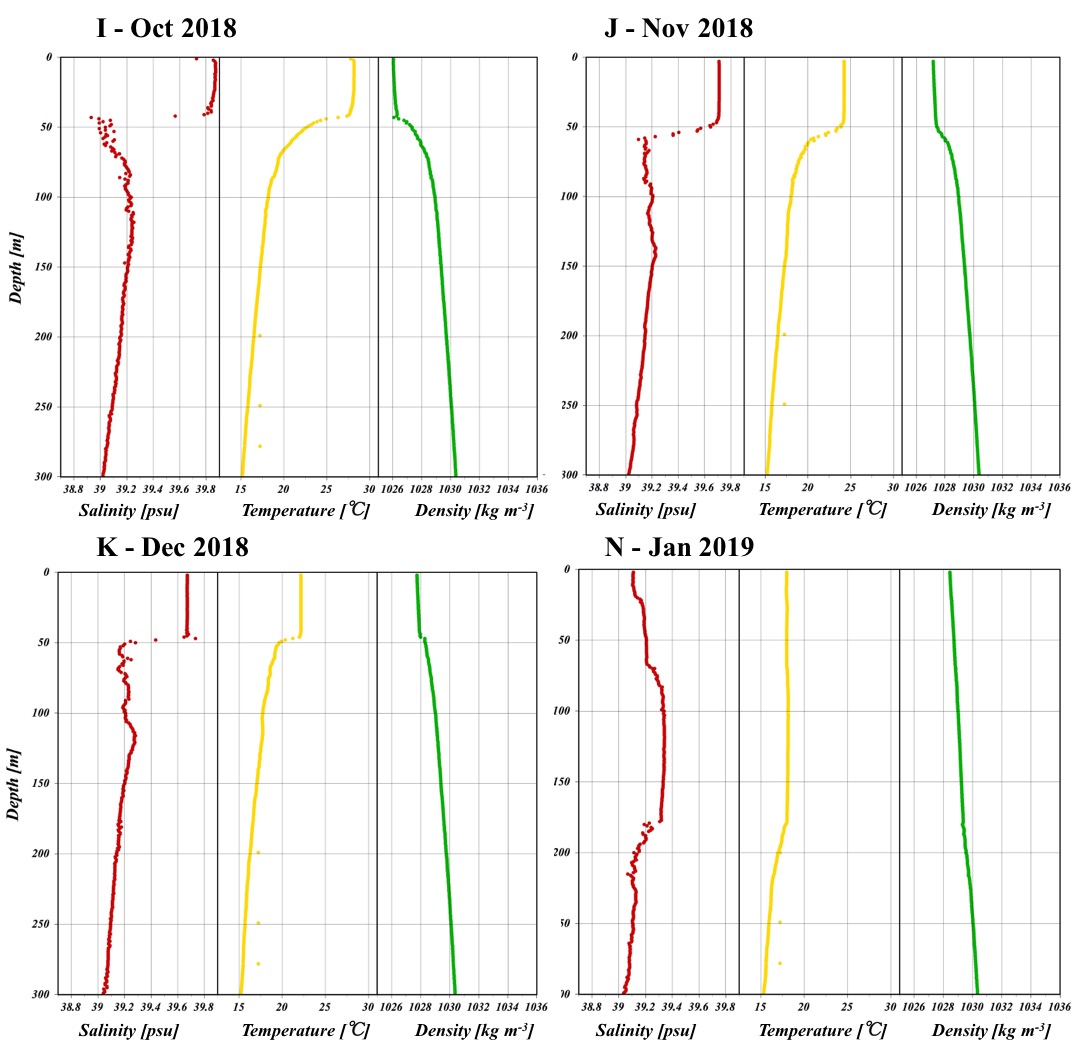

**Figure S1:** Temperature, Salinity and Density depth profiles (to 300m) for each of the 12 cruises to the THEMO-2 station. Note the clear pycnocline (“kink” in the density profiles) during May-December, which is not observed during February to April 2018 and January 2019. The MLD calculated when no pycnocline was observed ranged from ~21-186m. However, these values are misleading as they do not represent appropriately the actual mixing, which based on the relatively constant salinity gradient likely occurs much deeper. For this reason, the mixed period was defined as January-April, and no MLD is calculated for this period in Table 1.

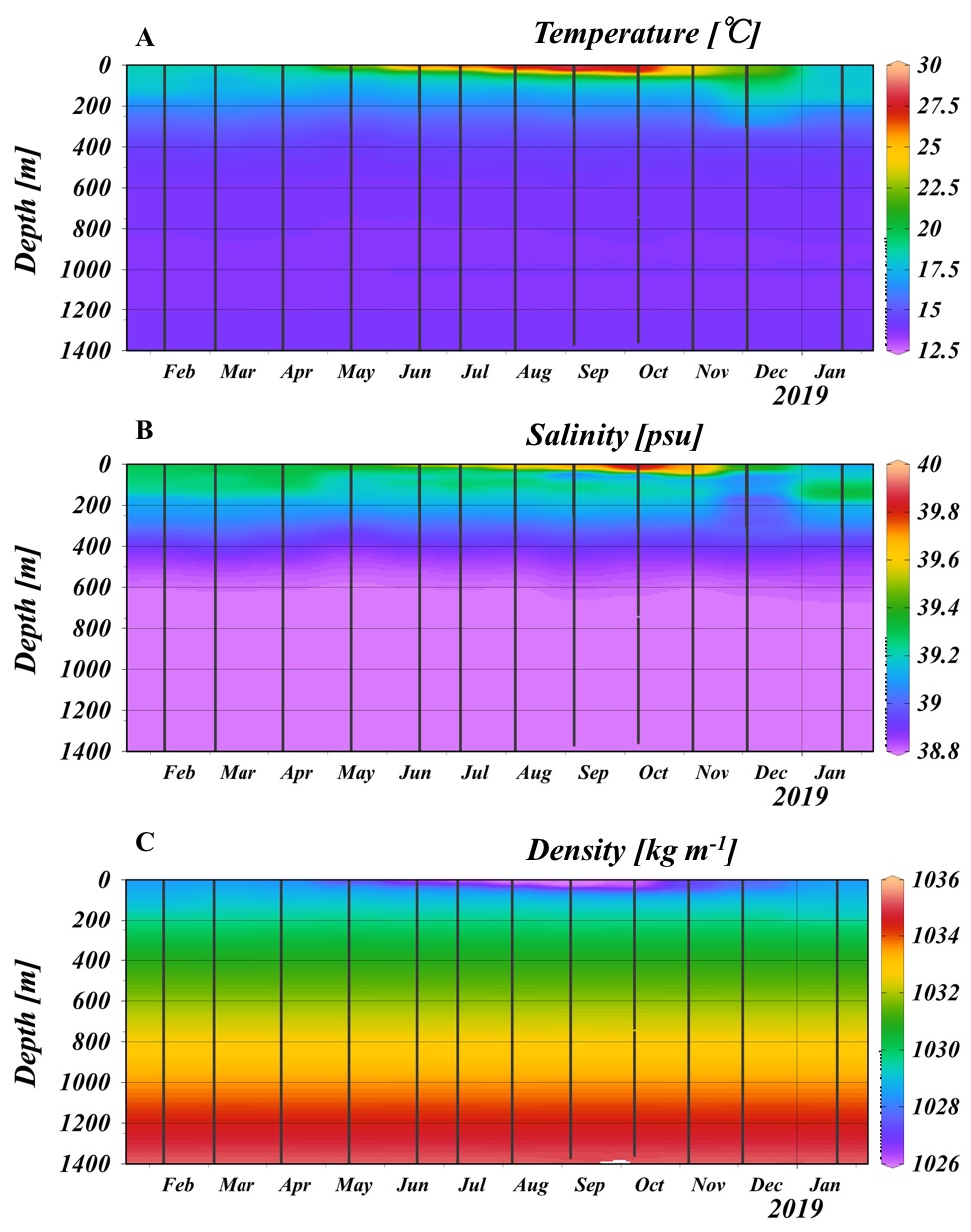

**Figure S2:** Physical parameters (temperature, salinity and density), across the entire water column at THEMO-2 (down to ~1,400m).

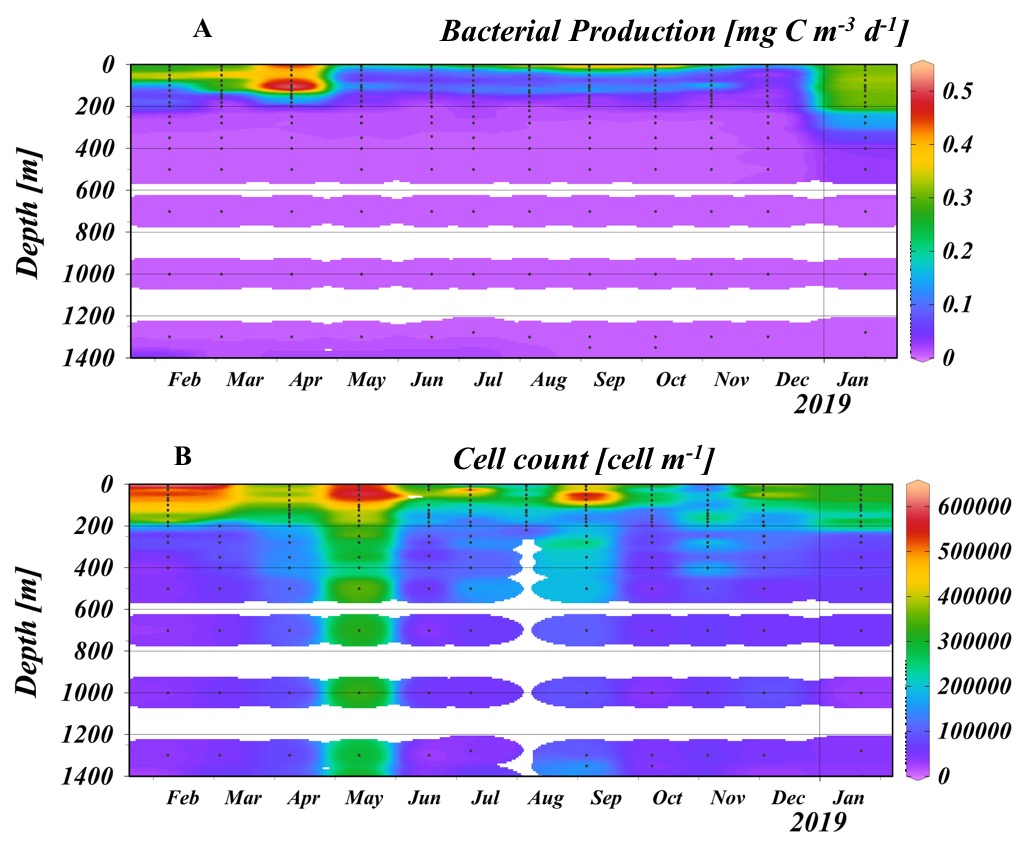

**Figure S3**: Production (A) and count (B) of total bacteria across the entire water column at THEMO-2 (~1,400m)

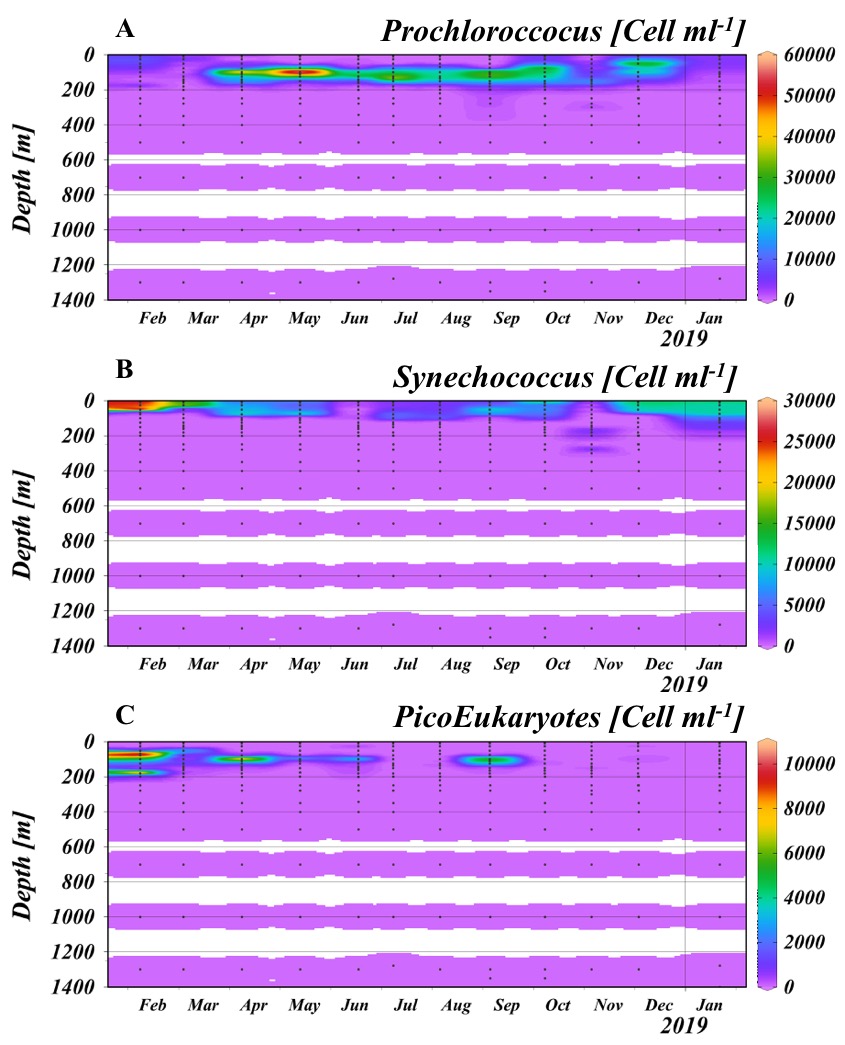

#### Figure S4: Cell counts of phytoplankton groups and of total cells across the entire water column at THEMO-2 (down to ~1,400m)

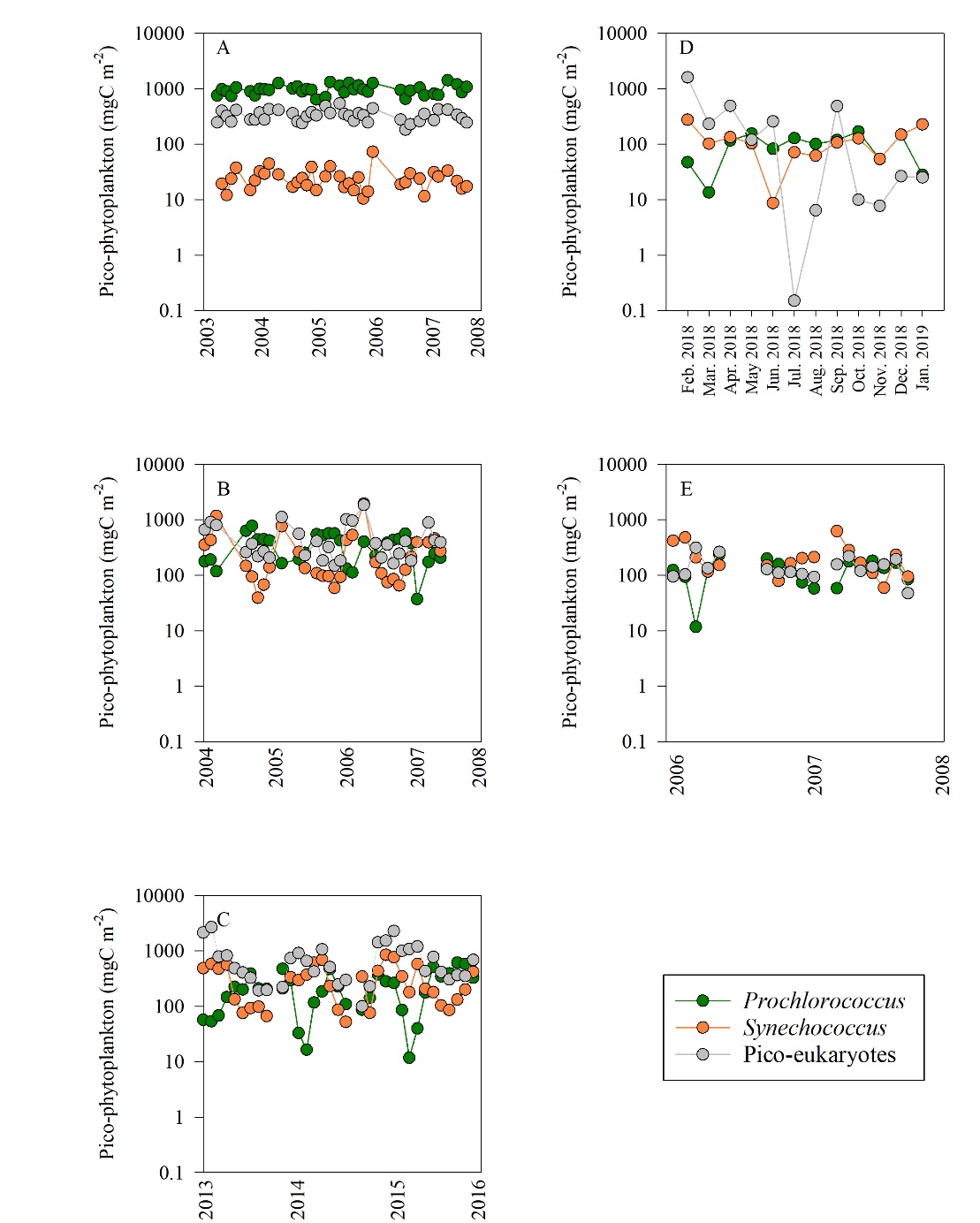

**Figure S5**: Integrated biomass time series of the calculated biomass of *Prochlorococcus* (green), *Synechococcus* (orange) and picoeukaryotes (gray) at stations HOT (A), BATS (B), the Northern Red Sea (C), and the EMS (D – this study and E - Yogev et al., 2011). Biomass measurements were calculated from the flow cytometry counts and assuming a *Prochlorococcus* biomass of 53 fgC cell^-1^, *Synechococcus* biomass of 175 fg C cell^-1^ and pico-eukaryotes biomass of 2100 fgC cell^-1^ (Campbell 2001). Data for stations HOT and BATS were compiled from (Malmstrom et al. 2010) and from the Northern Red Sea from the Eilat National Monitoring Program 2017.
